# Representational Learning from Healthy Multi-Tissue Human RNA-seq Data such that Latent Space Arithmetics Extracts Disease Modules

**DOI:** 10.1101/2023.10.03.560661

**Authors:** Hendrik A de Weerd, Dimitri Guala, Mika Gustafsson, Jane Synnergren, Jesper Tegnér, Zelmina Lubovac-Pilav, Rasmus Magnusson

## Abstract

Developing computational analyses of transcriptomic data has dramatically improved our understanding of complex multifactorial diseases. However, such approaches are limited to small sample sets of disease-affected material, thus being sensitive to statistical biases and noise. Here, we ask if a variational autoencoder (VAE) trained on large groups of healthy, human RNA-seq data of multiple tissues can capture the fundamental healthy gene regulation system such that the learned representation generalizes to account for unseen disease changes. To this end, we trained a multi-scale representation to encode cellular processes ranging from cell types to genegene interactions. Importantly, we found that the learned healthy representations could predict unseen gene expression changes from 25 independent disease datasets. We extracted and decoded disease-specific signals from the VAE latent space to dissect this finding. Interestingly, the gene modules corresponding to this signal contained more disease-specific genes than the respective differential expression analysis in 20 of 25 cases. Finally, we matched genes related to the disease signals to known drug targets. We could extract sets of known and potential pharmaceutical candidates from this analysis and demonstrate the utility in three use cases. In summary, our study showcases how data-driven representation learning using a VAE as a foundational model allows an arithmetic deconstruction of the latent space such that biological insights enable the dissection of disease mechanisms and drug targets. Our model is available at https://github.com/ddeweerd/VAE_Transcriptomics/.

## 2 Introduction

The human transcriptome reflects a network of genomic processes that control cellular functions. These processes are manifested as networks of interconnected groups of functionally related genes, and this modular topology has long been used to study fundamental cellular biology [1]. Various network disruptions, such as genetic changes and dysregulation, prevent them from operating correctly. While these disease factors may span the entire human interactome, they do not occur randomly but instead tend to co-localize in sets of functionally related genes, referred to as modules [2].

Studying such disease modules is essential for understanding pathophysiology, as holistic and mechanistic insights can be used to predict potential drug targets or biomarkers for disease prognosis [3]. Correspondingly, multiple computational methods are now available for understanding how disease changes co-occur in the gene regulatory network [4], with a substantial portion of work revolving around data mining. These data may, for example, measure gene expression [2], [5], [6]. Still, results have largely remained inconclusive and alternative benchmarks that go beyond GWAS validation have previously been conducted by us [6]. How to extract and interpret gene modules remains an open question.

The broader field of computational analysis of complex diseases has long struggled to find universally applicable methods that can distill knowledge from a broad selection of datasets. Complex diseases are inherently multifactorial, with many subtypes and differing characteristics. The amount of available transcriptomics data from healthy sources dwarfs that of most specific diseases [7], [8], and the significant advancement in our ability to model complex systems using large datasets with neural network models [9], has paved the way for a new era of computational analysis. Yet, studies constrained by limited sets of disease-related tissue material struggle to capture the essence of a disease change, and results often fail to generalize [10]. While transcriptomes reflect deep biology and disease-related perturbations, the limited number of disease samples and the challenge of identifying and using modules constitute a remaining hurdle slowing the discovery of novel biomarkers and drug candidates [11].

To what extent have recent advancements using neural networks been useful in mitigating the challenges associated with modules and a limited number of disease-relevant datasets? Indeed, deep autoencoders—neural networks designed to compress data to a significantly smaller set of variables—have been found to encode sets of functionally related genes using the same variables, known as hidden nodes, when applied to gene expression data [12]–[14]. However, these findings primarily suggest that autoencoders learn modularity to compress expression data rather than providing an approach to extract the most relevant latent structures for a specific disease state. In other words, how to extract and utilize the encoded structures remains an open question. Moreover, dissecting what input variables are encoded in which latent variable is not straightforward, despite recent progress in explanatory AI [15]. Furthermore, suppose the autoencoder is not variational in the latent space. In that case, the encodings in the compressed space do not necessarily have a continuously meaningful biological representation, leaving perturbations as a mean of model analysis flawed [16].

Inspired by foundational models in machine learning [17], we ask whether the learned representation from training a machine learning model using rich and extensive data from domain A, can be used to analyze data from domain B. In our setting, we explore whether a deep variational autoencoder (VAE) trained on healthy human RNA-seq gene expression profiles (rich data domain A) from the GTEx resource [7] can capture the fundamentals of human gene regulation. If so, can such a system-wide model be generalized to explain disease data (sparse data domain B)? Interestingly, we found that after training, the latent space of the VAE represented biological structures.

Moreover, we could extract hierarchical biological representations in the latent space, ranging from resolving cell types to identifying gene-gene interactions. Simultaneously, we observe that known disease features from genome-wide association studies (GWAS) are aggregated along the principal components of the latent space. Importantly, we also found this model to predict unseen gene expression data from The Cancer Genome Atlas (TCGA) [18]. These findings motivated us to design a generally applicable method for extracting gene sets that third parties with even small disease sample sizes of data can easily use. We then showcase this method on a compendium of 25 human disease datasets and found highly disease-relevant modules. Most gene sets exhibited a higher enrichment of associated disease genes than the top differentially expressed genes. This suggests that our approach identifies novel and intriguing disease-related genes beyond those with a clear disease signal in the data and yields more relevant disease related sets of genes, compared to the gold standard differential gene expression analysis of transcriptomic data. Finally, investigated disease gene sets showed significant enrichment of drug targets strongly associated with relevant pharmaceutical components for the analyzed diseases.

## 3 Results

First, a VAE was trained to compress human gene expression data to test the hypothesis that the learned representations in the latent space may embed biological processes. We continue by analyzing learned structures of the latent space and find biological patterns ranging from cell type to gene-gene interaction-specific structures. We show how elementary arithmetic operations on this latent parameterization of gene expression can be used to extract relevant disease modules when studying independent expression data from 25 diseases. The respective modules outperform differential gene expression analysis in predicting known disease genes in this analysis. Finally, we demonstrate that genes related to the parametrization of disease-related expression can be used to recommend pharmaceutical compounds, suggesting our approach to be important in drug repurposing studies (Fig. 1a).

**Figure 1:**
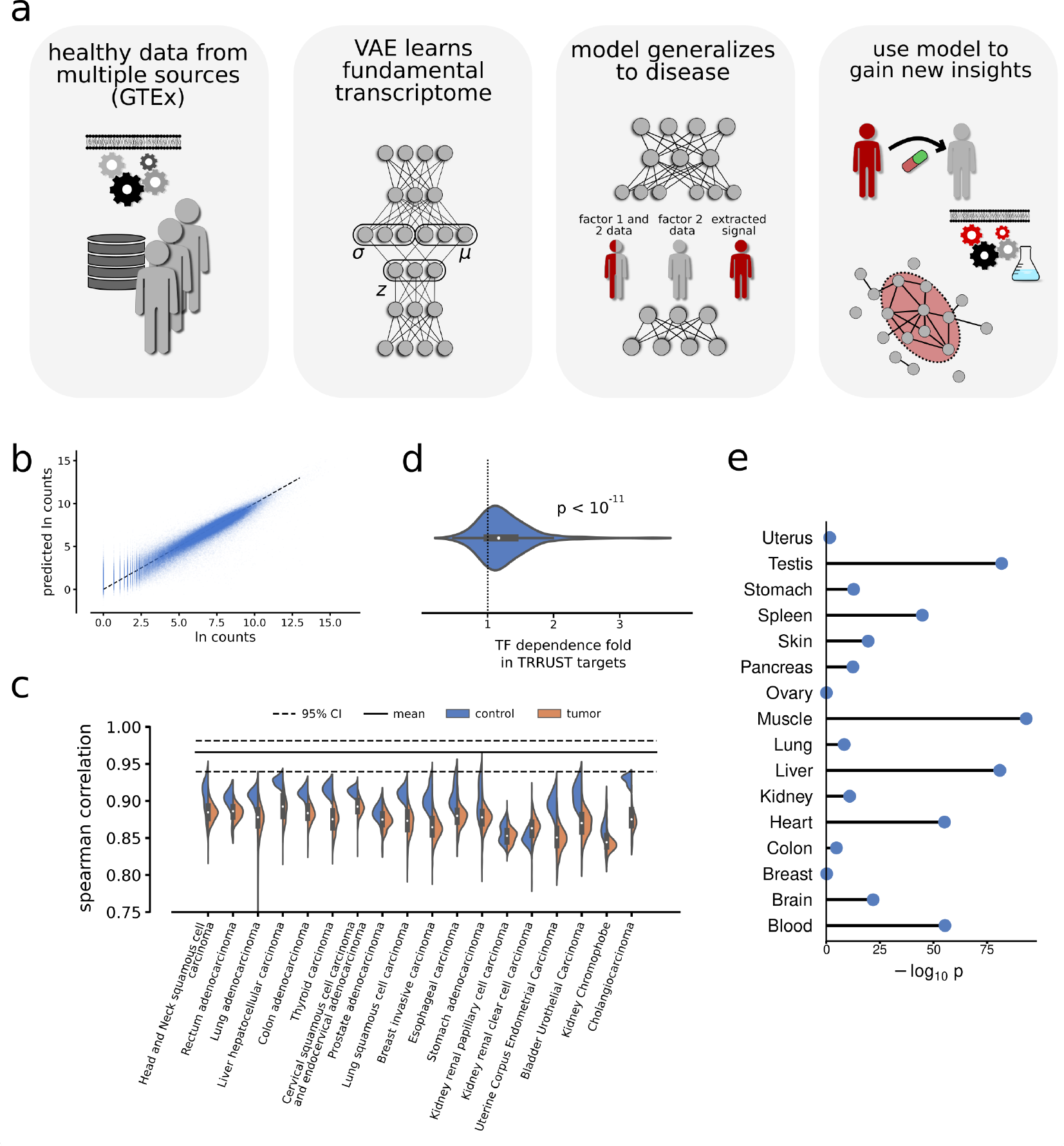
Model construction and validation. a) The workflow. The VAE model was implemented as a feedforward variational autoencoder. We train this model on uniquely healthy RNA-seq data and find it to generalize to disease data. Using simple arithmetic operations in the latent space, we extract specific disease vectors and use these vectors to extract disease modules and to suggest suitable drug candidates. b) The model performance on validation data is shown as blue dots. There was a clear similarity between the input and output of the VAE. c) The VAE was applied to independent data from TCGA, and the Spearman correlations over each transcriptomic profile were calculated. The figure shows the respective correlation of the GTEx data from b), plotted as a black line. In comparison, the independent data performed slightly worse, with healthy data (orange) performing slightly better than data from human tumors (blue). All correlations were highly significant. d) We tested the responsivity of known TF-target regulations by increasing a TF on the input level and measuring the changes on the output level of the VAE. The mean changes of known targets normed by the mean of all other genes on the output level were plotted, showing the distributions for the TFs. Most values were 1, indicating a stronger association in the model (binomial test p-value shown). e) We tested if cell types were enc1o7ded in the latent space and compared top-associated genes of 16 tissues in the data to tissue-gene associations in the TISSUES 2.0 database using a Fisher’s exact test. Shown are the*log*_10_ probabilities of the observed overlap or greater given H0: The sets are independent.

### 3.1 Training a variational autoencoder using healthy RNAseq profiles generalizes to unseen transcriptional disease profiles

The human transcriptome is a complex network of expressed genes that regulates numerous cellular functions and processes. To find a representation of this system, we designed a data-driven approach using a deep VAE to compress transcriptome data, with the rationale that the compression could encode biological structures into the latent space. We implemented the VAE (Fig. 1a) as a fully connected feed-forward neural network, with three hidden layers of 128, 2*64, and 128 nodes, respectively, where the middle variational layer contained 64 mean and standard deviation node pairs. Next, we used RNA-seq data from the 30 distinct healthy tissues in the Genotype-Tissue Expression project (GTEx), randomly divided into 11,258 training, 4,362 test, and 1,762 validation samples unused in model training. We evaluated the data loss of the trained model after compression using the validation data estimated with the Spearman’s rank correlation for each validation expression profile. We found a mean correlation of 0.96 (95% confidence interval = 0.94-0.98) (Fig. 1b-c).

Knowing that the model could faithfully recreate healthy human material, we next aimed to validate the learned compression on independent disease data. To this end, we used case-control datasets from The Cancer Genome Atlas as processed by Wang et al. [19], noting that cancer is one of the most heavily perturbed disease systems. We ran each tissue and respective tumor gene expression profile through the trained VAE and calculated the Spearman’s rank correlation between model input and output for each transcriptomic profile. Strikingly, we found strong correlations before and after compression for all datasets (mean Spearman’s *ρ* = 0.87, Fig. 1c), with the healthy controls (mean *ρ* = 0.90) performing slightly better than the tumor data (mean *ρ* = 0.87). In other words, the variational autoencoder that had learned to compress expression data of healthy material could be used to represent and compress unseen transcriptional disease profiles. In comparison, in a previous study, we built a feed-forward neural network to predict transcriptomic profiles from transcription factor expression, where we reported an out-of-sample prediction on tumor data with 80% predicted variance. [9].

### 3.2 Rich biological multi-scale representation in the learned latent space

During the training of an autoencoder, data structures of varying resolutions are known to be encoded into the latent space [17]. The data structures, although nontrivial to extract, are exciting since they potentially hold biological insights unbiased by *a priori* biological assumptions. We, therefore, aimed to examine these biological structures further.

The training data of the variational autoencoder contained 30 distinct healthy human tissues divided into 54 subcategories, and we hypothesize that the differences between tissue types are the most significant source of variation in the training data. We therefore examined whether tissue-specific gene sets could be extracted from the latent space. To this end, we downloaded the knowledge channel of the TISSUES 2.0 database, containing manually curated known associations between gene expression and human tissue types [20]. We matched the 30 distinct tissue types in our data to the tissue types present in TISSUES 2.0, considering only literal matches, i.e., “Muscle” to “Muscle”, and so forth. Next, we filtered the matches further by requiring at least 100 tissue-gene associations in the database and at least 10 samples for each tissue type in our test set, resulting in 16 distinct tissue types.

To extract tissue-specific gene sets, we employed the strategy described in the Methods section (specifically Eq. 1-4). We analyzed the top 500 ranked genes for each of the 16 tissue types for tissue-specific enrichment using a Fisher’s Exact test. Notably, we found major enrichments for 14 of the 16 tested tissue types, with exceptionally high enrichments for the Heart, Liver, Muscle, and Testis (Fig. 1e).

We next analyzed biological pathways to assess if more fine-grained resolution cell structures were encoded in the latent space. Since there is arguably no approach to directly extract learned features from the latent space directly and to analyze the latent structures, we decided to perform a linear principal component analysis on the compressed validation data from the GTEx dataset. We then augmented the latent space compression of the data by increasing the activations along each principal component (Methods). Next, we considered the top 1,000 changing genes for each principal component perturbation and calculated the overlaps with genes associated with KEGG pathways. We found 10,684 statistically significant principal component-KEGG pathway overlaps (median odds ratio, OR = 38.5, 50% on the interval [21.2, 77.2]) using a Bonferroni-corrected Fisher’s exact test. This suggests that biological processes are represented in the latent space.

These results motivated us to test the latent space for even more fine-grained interactions, and we continued to study individual gene-gene interactions. We have previously shown how neural networks can use transcription factors (TFs) to predict target gene expression [9], and we herein used the same approach to associate TFs with target genes. For each TF, we increased the expression in the validation data with 5 standard devia-tions and measured the gene response on the output level, defined as the absolute gene change from the TF increase. We measured the mean of this response for known target genes in the TRRUST [21] database and found on average 24% higher levels than of the genes not registered as known target genes in TRRUST, with 140 of 190 TFs having target genes with higher variance (binomial test p *<* 2.4∗10^*-*11^, Fig. 1d). Even though the increase was lower than the 70% found in our earlier work [9], we note that model to have been specifically designed to capture TF-target regulations.

### 3.3 Known disease genes aggregated along different principal components in the latent space

With results indicating that biological mechanisms from the healthy data were incorporated into the latent space structure, we also sought to test if the same was true for disease mechanisms. We, therefore, downloaded The GWAS Catalog [22] and again augmented the principal components of the compressed latent space using the validation data. For each principal component, we calculated the overlap of the top 1,000 responding genes with genetic variations associated with disease in the GWAS catalog. Using a Bonferroni correction for multiple testing, we found 311 significant disease-principal component pairs. Moreover, we found 4,515 nominally significant pairs, a 5.3fold increase of what would be expected under the null hypothesis of random overlap (expected 857.6 out of 17,152 at 0.05 rejection rate).

### 3.4 Algebraic manipulation of identified case-control vectors in the latent space extracts highly relevant disease modules for 25 diseases

A core advantage of the VAE is the inherent continuous encoding of representations in the latent space. This representation of data structures has allowed earlier work in computer vision to show how images can be manipulated along an axis that outputs a continuous change in an encoded feature [23]. Here, we explore whether an analogous operation could be translated using gene expression data. Since disease mechanisms were captured in the latent space structure, we decided to search for a direct approach to extract clusters enriched with known disease genes from the latent space. To this end, we downloaded all relevant case-control studies of complex diseases from the Expression Atlas database (see Methods for inclusion criteria).

Moreover, we also included five case-control studies used in our latest study on gene module inference [24], bringing the total number of disease sets to 25. We hypothesized that the difference between the two conditions could be manifested as a vector in the latent space activations. In other words, for each disease, we searched for a vector in the latent space that corresponded to the respective transcriptomic changes. To derive such latent vectors, we compressed the case and control samples separately, and for each node in the latent space, we calculated the average difference between the two conditions. We next amplified this disease signal encoded in the latent vector by increasing the activation values threefold. Notably, the inherent VAE property of continuous representation of features in the latent space enables such an amplification to translate into information on disease-relative gene changes after decompression. To identify the impact of the amplification of the disease-relevant vector in the latent space, we decompressed the now disease-augmented data and, as a comparison, generated 1,000 random, normally distributed latent space variables and subsequently decompressed these profiles to the complete transcriptomic profiles. Next, we computed the rank for each gene in the decompressed latent disease vector and compared it to the 1,000 random profiles gene ranks for reference. The method is further described in the Methods section, and a schematic representation can be found in Fig. 2a. Gene modules are often defined as sets of genes that share a biological function and, therefore, are densely connected on a gene interaction graph. To analyze the modularity of the output genes of our method, we applied it to our compendium of 25 disease expression sets. We mapped the top 500 output genes to the STRING protein-protein interaction (PPI) network and calculated the edge enrichment for each gene set using the STRINGdb R package [25]. All gene sets were highly modular regarding increased edges between the genes within each module (median fold of expected edges: 6.1, Fig. 2b). We next filtered our generated gene sets to only contain the largest component connected in the STRING PPI.

**Figure 2:**
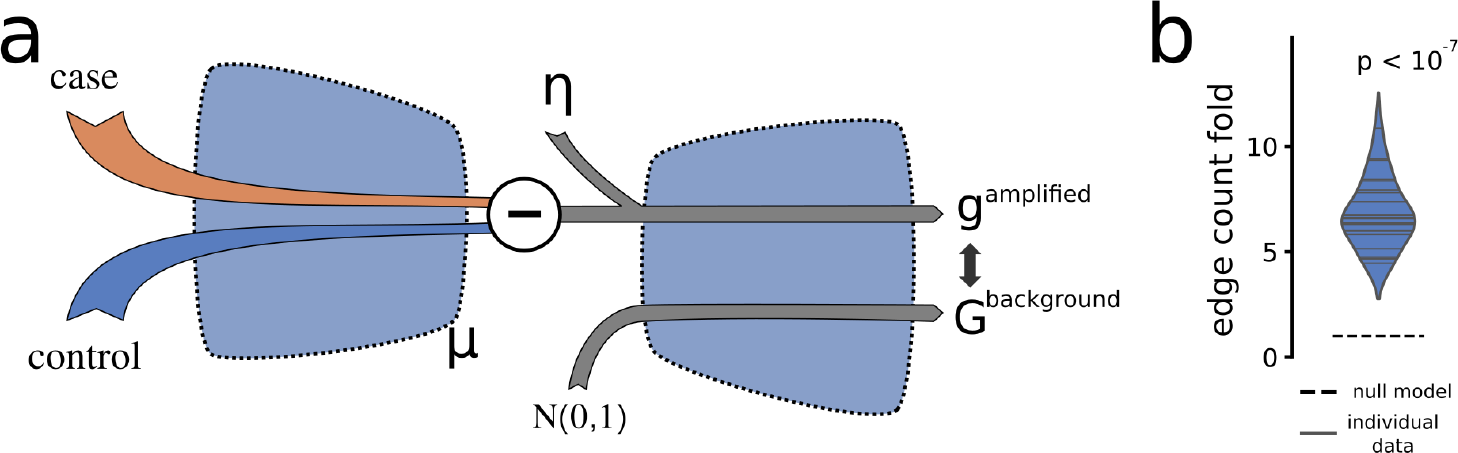
Highly modular gene sets could be extracted. a) We devised an approach to extract relevant groups of genes from factorial data, i.e., data containing a disease case and control samples. These two factors were compressed to the latent space, and the respective mean difference was calculated, denoted by the disease vector. Next, the disease vector signal was amplified with a factor *η* and decompressed. The decompressed values were then compared to decompressed random, normally distributed data. b) For the 25 studied disease datasets, we used our algorithm to extract sets of genes and used the STRING PPI network to calculate the enrichment of known interactions between these genes compared to the expected number of interactions. All gene sets had highly significant increases, with the increase typically being 5-fold or more. The p-value shows the probability of all 25 samples being greater than the null model.

### 3.5 Extracted modules are more enriched with disease-related genes than the respective observed Differentially Expressed Genes

Knowing that the output gene sets were highly modular, we henceforth considered the sets as gene modules. We next compared the modules to the corresponding known disease-associated gene sets in the DisGeNET database. We found clear enrichments of disease-associated genes across next-to-all diseases, including genes that were not found to be differentially expressed in the data source (Fig. 3a). For each disease module, we again performed a Fisher’s exact test for enrichment of known associated disease genes and found striking enrichments of overlapping genes, typically having OR between 4 and 8 (Fig. 3b).

**Figure 3:**
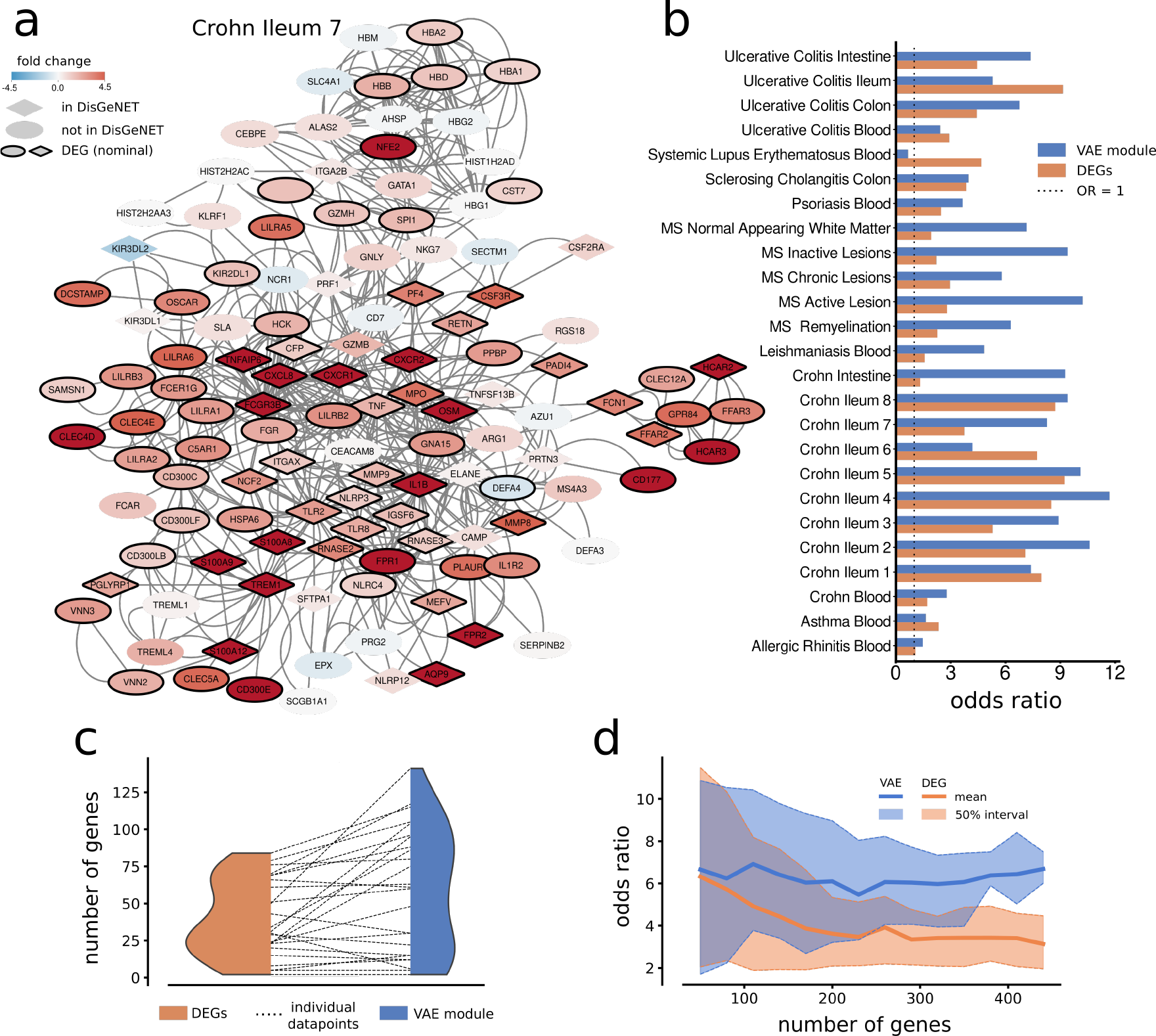
Modules were significantly enriched for known disease genes. a) An example of a module of size 100 genes, extracted using RNA-seq data from samples of the ileum of Crohn’s disease patients and healthy controls. As can be seen, there are several genes that were not differentially expressed in the data but associated with Crohn’s in DisGeNET. In b), the OR of Fisher’s exact test for overlap with DisGeNET genes is shown for the VAE module and the respective top DEGs. The modules were extracted with a size of 300 genes, which is the default setting. Strikingly, the VAE-derived genes are more relevant for each disease than the corresponding top DEGs in the data. c) The distribution of DisGeNET genes in the top DEGs and in the equally sized respective modules are shown for the diseases. The increase in the number of genes for each disease is shown with the dotted line between the two distributions. d) The mean overlap of the VAE modules and top DEGs with the DisGeNET genes for all diseases, as a function of the number of genes, showed a better performance for the VAE modules. Interestingly, the modules seemed to be stable in terms of OR with respect to module size, suggesting our approach to be robust.

We next compared the number of disease-associated genes from the DisGeNET in the disease modules with the occurrence of DisGeNET genes among top differentially expressed genes (DEGs) and found the number of genes to be larger in the modules in most diseases (Fig. 3c). Specifically, we used the DEG analyses accompanying the expression data from the Expression Atlas, and for the separate multiple sclerosis (MS) datasets we used the DEG analysis provided by Elkjear et al [26]. First, we filtered the DEGs to contain only genes in the STRING database and removed the DisGeNET disease-associated genes not present in the DEG data to ensure genes were drawn from the same pool as the modules. Moreover, we used a Fisher’s Exact test between the equally sized set of top DEGs and the corresponding DisGeNET disease-associated gene sets to test for enrichment. In 20 out of 25 cases (Fig. 3b), the modules derived from the latent space using our trained model had a higher OR of overlap enrichment than the corresponding number of DEGs. (binomial test 20 of 25: p *<* 0.002). We also analyzed the impact of varying model sizes from 50 to 450 included genes and found the performance of our method to be surprisingly robust to module size. Furthermore, we found the modules of varying sizes to consistently outperform the DEG analysis in terms of DisGeNET overlap (Fig. 3d). In other words, the gene modules extracted using our VAE were a more relevant foundation for disease analysis of factorial data than the gold standard technique; the DEG analysis

### 3.6 Enhancing the Disease vectors suggests suitable pharmaceutical compounds

Having studied how the arithmetic extraction of disease-associated vectors in the latent space carried information on disease-associated genes, which arguably allows for further studies of pathophysiology and pathogeneses, we next tested how this parametrization could predict suitable pharmaceutical compounds. Analogous to our study of GWAS genes, we analyzed the top 1,000 genes associated with each principal component of the latent space. We then iteratively compared each set of 1,000 genes to the known targets of each drug with more than 10 known target genes in the DrugBank database.Specifically, for each of the 64 sets of 1,000 genes, we performed 264 Fisher’s exact tests to enrich drug targets. We found 62 of 64 gene sets to have more significant associations than what would be expected by random (expected 32, p_*binom*_ *<* 1.2∗10^*-*16^.

We continued the analysis by matching the top genes in the decompressed output of the respective disease vectors to the known targets of the pharmaceutical compounds in DrugBank. In other words, we could extract potential pharmaceutical compounds to aid drug repurposing by associating a vector that represents non-linear disease structures in the latent space to pharmaceutical compounds via the VAE decoder. Importantly, the top-ranking drug suggestions for each disease were strikingly relevant. Here, we present three cases, while the ranked pharmaceutical compounds for all tested diseases are found in Supplementary material I.

**Case 1**; Multiple Sclerosis. For the MS data presented in [26], labelled MS active lesion in Fig. 3, the top drug suggestions were Muromonab, previously tested for MS [27], Ibrutinib, a Bruton’s tyrosine kinase inhibitor (BTKi), which are all highly relevant in MS research [28], The third-ranked drug was Daclizumab, which has been approved as a treatment for MS [29], but retracted due to adverse side effects. The fourth was Zanubrutinib, another BTKi compound, and the fifth was found to be Alemtuzumab, an already used treatment for MS.

**Case 2**; Crohn’s disease. We next tested the approach on Crohn’s disease, labelled “Crohn Ilium 1” in Fig. 3 [30]. The top two matching compounds were zinc and zinc acetate, and zinc deficiency has been associated with poor clinical outcomes in people with inflammatory bowel diseases [31]. The third compound was Dilmapimod, a p38 mitogen-activated protein kinase (MAPK) inhibitor with known anti-inflammatory effects [32], the fourth was Glucosamine, which has been used in cases with osteoarthritis, while the fifth was VX-702, another p38 MAPK inhibitor.

**Case 3**; Systemic Lupus Erythematosus (SLE). Analyzing the data in [33], labeled Systemic Lupus Erythematosus (Blood) in Fig. 3, we found the top predicted drug to be Gemcitabine. This chemotherapeutic agent has been reported to, albeit rarely, trigger SLE [34]. We note that our approach does not consider the direction of gene dysregulation. In fact, in the top five predicted compounds, three are primary used as chemotherapeutic agents (Gemcitabine, Enzastaurin, and Sunitinib), which we speculate to be related to the anti-nuclear immunological response of SLE. The remaining two drugs were Fostamatinib, a tyrosine kinase inhibitor used in immune thrombocytopenia, and Cladribine, selective lymphocyte depleting agent that is primarily used in the treatment of MS. We note, however, that Cladribine has been tested as an agent against SLE [35]

### 3.7 Underlying structures in the latent space restrict the modes of analysis

The analyses we present herein suggested that arithmetic operations in the latent space of a VAE trained on biological data can be used to extract meaningful biological structures. Lastly, we asked how the vector amplitude in the latent space translates to meaningful biological interpretations on the decoded level. Starting with the analysis of disease module prediction based on case-control data (Fig. 3), we hypothesized the differences between case and control samples to indicate disease signal and, in extension, method performance. We thus calculated the mean Euclidean distance between the compressed case and control vectors in the latent space. We found this metric well-correlated with the previously presented odds ratios of disease gene enrichments (Fig. 4a, Spearman’s *ρ* = 0.52, *p <* 0.008).

**Figure 4:**
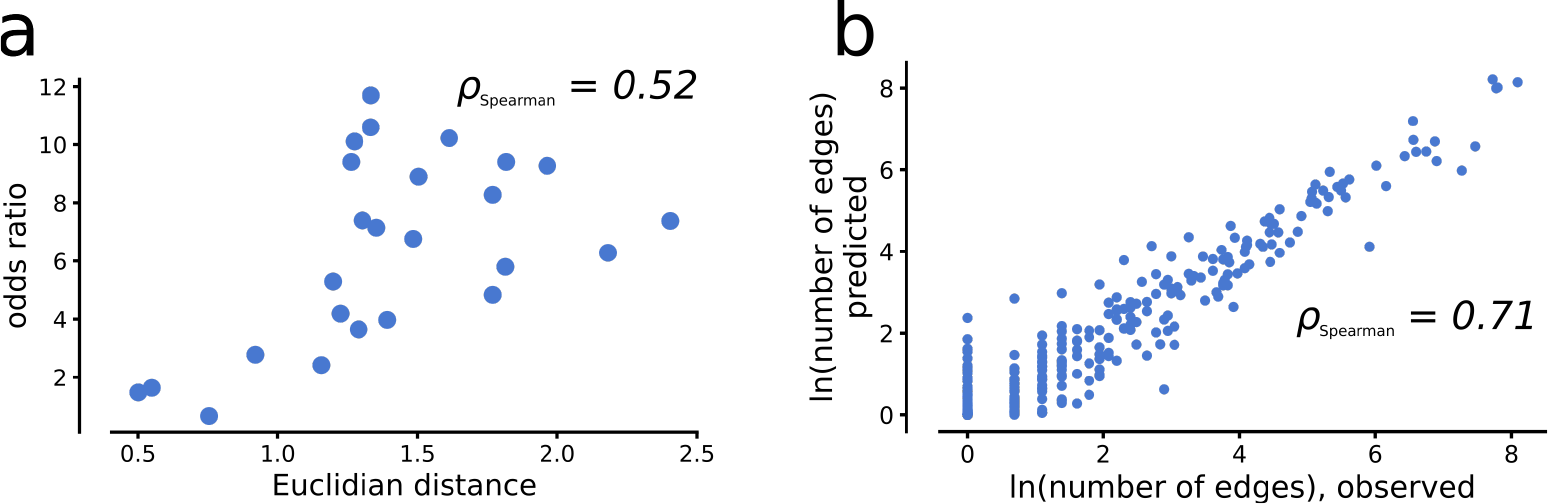
High performance is connected to defined areas of the latent space. a) We found the Euclidean distance between the case and control vectors in the latent space to be well correlated with the observed enrichment of known disease genes on the output level. b) We trained a regressor model to predict the number of edges on a PPI network based on random vectors in the latent space and found a clear relationship.

We asked if vectors in the latent space correspond to an interpretable set of genes, and to this end, we generated 100.000 random latent space profiles. Typically, the vectors in the latent space that capture transcriptomic changes between disease and healthy state have, on average, low activation values compared to vectors of a compressed whole transcriptome. We believe this change in amplitude to originate in the disease changes to be a subpart of the broader transcriptome signal. To mimic these low activation values, we sampled the activation values from a standard normal distribution with *µ* = 0 and *σ* to be a random number from the uniform distribution between 0.01 and 0.4. Next, we translated the random latent space profiles into a module, as described under “Methods–module extraction”, but without the augmentation of the latent space activations and calculated the number of edges in the output module. We found most sets to have zero or few PPI interactions between them. Furthermore, we trained a regressor model to predict the number of PPI interactions (Methods). We found a precise performance on a separate test set of 10.000 profiles prepared similarly (Fig. 4b), suggesting the modularity to be encoded within a well-defined area of the latent space.

## 4 Discussion

Here, we have presented how a VAE can learn to represent the fundamentals of transcriptomics associated with human tissues. We show this data-driven approach to generalize from exclusively healthy material into disease-affected domains, making representation learning practical. Moreover, we demonstrate how to extract biologically relevant insights from the latent space, thus being of broader value for biomedical research. We showed how the latent vectorization of gene expression data can be utilized such that arithmetic functions can extract important disease signals. Using data from 25 independent disease datasets, we extracted sets of disease-relevant genes, herein referred to as disease modules, and compared them to previously known disease genes. All but one of the modules were significantly enriched for known disease genes, and notably, 20 out of 25 cases outperformed the corresponding top differentially expressed genes. Furthermore, we show how these modules could be used to predict drug candidates with high therapeutic potential from approved and registered pharmaceuticals.

We hypothesize that the model’s performance originates from the encoded biological structures embedded in the latent space during training. While the VAE was trained using exclusively healthy human gene expression profiles, the model could still explain previously unseen disease mechanisms. Importantly, we do not claim that our VAE has learned all cellular mechanisms. Neither is there a fully established approach to extracting the encodings of the latent space, and our analyses of these mechanisms, i.e. the cell type, pathway, and TF binding analyses, are limited. However, we claim that the model represents important transcriptome processes such that the model can be used to understand at least a substantial set of human diseases.

Nevertheless, the model could potentially be improved. The GTEx data we used is limited to *<*20,000 samples; arguably, most variation stems from different cell types. We also note how others have found cell type to be the predominant parameter for latent space representation [36]. While there are several other sets of massive gene expression datasets [7], [37], we believe using the GTEx consortium data is a key strength of our analysis when applied to independent disease data. Furthermore, we note that the gold standard, i.e., the DisGeNET, used herein, is not a complete set of relevant disease genes. Instead, all genes taking part in the generated modules are of interest from a mechanistic and a biomarker perspective.

This approach is part of a larger paradigm shift, where the availability of big biological data coupled with the potential of deep learning allows for complex biological features to be modeled without relying on what most often are greatly limited datasets. Way et al. (2020) [13] and Dwivedi et al. (2020) [12] have recently shown autoencoders applied to gene expression data to learn cell functionalities, incrementally increasing with model depth. This work contrasts Way et al. and Dwivedi et al., both used independent machine learning models to extract specific encoded structures from latent layers, in that, our method is generally applicable with the end goal of being a simple tool available for users. In Seninge et al. (2021) [38], a VAE was built to compress gene expression data to study gene modularity. The model was constructed with a linear decoder, with modules instead being supplied by the user to study cell regulation, as opposed to our work, where the gene modules are the desired output. An exciting work by Kuenzi et al. (2020) [39] describes the DrugCell model, in which gene modularity was hard-coded into a neural network to predict drug outcomes in an interpretable latent space. While these studies used gene modularity in their modeling approaches, we note that prior knowledge is prone to bias.

Yet, it should be noted that the broader challenge of explainable AI has caught significant attention recently. There have been numerous efforts to design encoder-decoder architectures such that the representations in latent space can be interpreted [40]. This has proven to be difficult, and, in our view, previous attempts in the machine learning community have been too general and not considered the specifics of the molecular data. This remains a key challenge when using machine learning in a biological context [41], [42]. Our present work and others have shown a way forward in addressing this important challenge of understanding what machine learning models learn when trained on biological data. This challenge is now renewed in the recent context of the emergence of foundational models in bioscience based on single-cell genomics data [43].

Our study is, in this sense, part of a growing approach to machine learning in life sciences that aims to capture the fundamentals of a system to make explainable predictions that generalize outside of training data accurately. Theodoris et al. (2023) [44] used 30M single-cell transcriptomic profiles to understand the fundamentals of network biology and transferred these understanding features to context-specific predictions. Dalla-Torre et al. (2023) [45] show how a foundational model of human nucleotides could be used to predict molecular phenotypes. In a notable work by Lotfollahi et al. (2023) [46], single-cell data were mapped using a nonlinear encoder paired with a linear decoder and a biologically informed latent space to interpret gene processes. While Lotfollahi et al. presented intriguing results, we consider an entirely data-driven model learning, yielding explanatory predictions, to be a strength of our method.

The main scientific contribution of this study is the novel applicability of VAEs to extract useful information from the latent space representation and thereby enable an analysis of biological signals in human RNA-seq experiments. The model, together with the accompanying Python code, is freely available to users (Availability), and we believe the software and the scientific insights will serve as a stepping stone towards moving analyses of high-throughput data into the AI era. Whereas our study focused exclusively on gene expression, extracting and studying cellular mechanisms in the latent space of a VAE is also applicable to other data, such as methylation, phosphorylation, and protein abundance data. We show how arithmetic operations on the latent space of a VAE can be used to extract and enhance disease signals, which we showcase by studying these disease signals from a disease module perspective. Furthermore, we also show how these vectors can be used to suggest candidate pharmaceutical compounds, which is an end goal of bioinformatic analyses. Yet, these approaches only showcase the model approach from a directly applied disease perspective, and we speculate that future applications can make use of the generative feature of VAEs in general, coupled with our suggested arithmetic operations related to disease signals to study disease mechanics and offer possible treatments in novel, informative ways.

## 5 Methods

### 5.1 Data extraction and normalization

The Genotype-Tissue Expression project (GTEx) was used as training data. We downloaded GTEx Analysis V8 with the dbGaP Accession phs000424.v8.p2, containing 17,381 healthy human RNA-seq profiles. We randomly partitioned the data into training (65%), test (25%), and validation sets (10%). To filter out less relevant genes, we downloaded the STRING human protein-protein interaction network [47] and removed all genes not included in the STRING network. We normalized the gene raw count values with only a logarithmic transformation, a technique we and others have advocated [9], [36]. The Cancer Genome Atlas gene counts were downloaded in a preprocessed form from [19] and processed in the same way as the GTEx data. For the GWAS data, we downloaded the “All studies” v1.0.2 from the GWAS catalog and used the “REPORTED GENE(S)” column for the gene annotations. Moreover, we removed all traits with less than 100 associated genes. For the disease-control gene expression data we searched the Expression Atlas with the following inclusion criteria: The organism should be Homo sapiens, the technology should be RNA-Seq mRNA, The type of experiment should be differential, it should be a case-control experiment, raw counts should be available, the sample tissue should be in our training data and, for validation purposes, as implemented in Dviwedi et al 2020 [12], there should be at least 100 known gene-disease associations present in the DisGeNET database. Moreover, we recently studied [24] how upstream regulators of modules can be inferred using the data published by Elkjaer et al. (2019) [26] and also included this case-control dataset in the analysis.

### 5.2 Model design and training

We implemented the model in the Python package Keras using a feed-forward neural network structure. The encoder was built with one input node per each of the 16,819 genes in the data and a dropout rate of 20%. The encoder hidden layers had 128 and 64*2 hidden nodes, respectively, where the 64*2 encompasses a mean and a log-transformed variation. The decoder was built to sample these mean and variation nodes and decompress the data to the full gene set size. All nodes were implemented with the leaky ReLU activation function. The model is available for download at https://github.com/ddeweerd/VAETranscriptomics/.

The initial training consisted of 500 epochs with the reconstruction error multiplied by 100 while keeping the Kullback–Leibler (KL) divergence error constant. Subsequent training consisted of 100 epochs, each with diminishing reconstruction error multiplication factors of 50, 20, and 10 until the KL divergence and reconstruction error were in equilibrium.

### 5.3 Latent space analysis

We analyzed if and how KEGG pathways were represented in the latent space. To this end, we used the validation set from the GTEx data and compressed it to the latent space. We used a linear principal component analysis to extract vectors of node activation combinations and increased the node activations 5 times along each vector independently. Measuring the decompressed changes, we selected the 1,000 top changing genes of each vector and used a Fisher’s exact test to enrich genes in KEGG pathways.

### 5.4 Module extraction

Our module extraction method was implemented in two steps. First, we identified a vector corresponding to the difference between two states in the latent space. Next, we decompressed this vector to find its primary encoded gene expression variables.

The latent space vectors were derived from the latent space by compressing the case and control samples separately. For each node in the latent space, we calculated the average difference between the two factors. In detail, the mean activation vector **z** was calculated as the mean values of the *µ*-nodes inherent to a VAE. The vector **z** was calculated for each set, and the difference between disease and control activation vectors ***ν***_*case-control*_ was calculated (1).

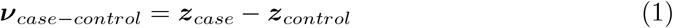

Next, the vector ***ν***_*case-control*_ was increased with a scalar factor *η*, with a default value of 3, and decompressed using the decoder *f*, as expressed in (2). In (2), ***g***denotes the decompressed gene expression vector from the compressed disease vector ***ν***_*case-control*_. Notably, the inherent VAE property of continuous representation of features in the latent space is a prerequisite for this amplification to translate well into information on disease relative gene changes after decompression.

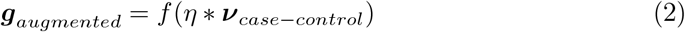

To analyze ***g***with respect to the normal gene expression background, a matrix of 1,000 normally distributed random latent space variables, denoted by ***X***, was decompressed to the background gene expression profile ***G***_*background*_, (3). We remark that ***G***_*background*_ is a matrix containing 1,000 random gene expression profiles.

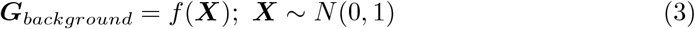

Lastly, to estimate the most relevant genes in the vector ***g***; we ranked each gene *i* for the number of times where the *i*:th gene, *g*^*i*^, was larger than the corresponding *j*:th random sample for gene *i*. In other words, we counted the number of times the inequality in (4) was satisfied, and extracted the top ranking genes for further study.

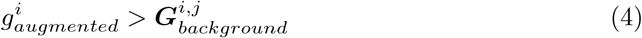

To filter genes when compared to the STRING PPI network with a confidence score cut-off of 700, we extracted the largest connected component subset of the genes and considered this set as the extracted disease module.

Since our study of tissue-specific genes had no case-control data, we generated 1,000 normally distributed random latent space variables and decompressed them to full transcriptomic profiles. The decompressed random samples were used as a control group, and the total number of samples in our GTEx test set for each distinct tissue type served as the case samples. Then, both the case and control samples were compressed to the latent space, and the average activation of the control samples was subtracted from the average activation of the case samples, resulting in a latent space vector that, we hypothesized, encodes the difference between the two conditions. We decompressed this vector to a transcriptomic profile to investigate if relevant gene sets were captured in the latent space vector. Furthermore, we sampled an additional 1,000 normally distributed random latent space variables and decompressed them to full profiles for comparison.

## 6 Acknowledgements

This work was supported by the Systems Biology Research Centre at the University of Skövde under grants from the Knowledge Foundation (grant no. 20200014 R.M, Z.L.P, J.S), the Assar Gabrielssons Fond (grant FB21-66, R.M, H.D.W), the Swedish Research Council (grant no. 2019-04193, H.D.W, M.G.). The computations were enabled by resources provided by the Swedish National Infrastructure for Computing (SNIC), partially funded by the Swedish Research Council through grant agreement no. 2018-05973.

## 7 Availability

The model, accompanied by relevant Python and R code is available at https://github.com/ddeweerd/VAETranscriptomics/.

## Notes

https://github.com/ddeweerd/VAE_Transcriptomics/

### Competing Interest Statement

The authors have declared no competing interest.

